# High pathogenicity avian influenza virus H5N1 clade 2.3.4.4b in Antarctica: Multiple Introductions and the First Confirmed Infection of Ice-Dependent Seals

**DOI:** 10.64898/2026.01.04.697571

**Authors:** Jane L. Younger, Laura Patier, Talia Brav-Cubitt, André van Tonder, Augustin Clessin, Jamie Coleman, Jessica Farrer, Clive R. McMahon, Amandine Gamble, Nicholas Fountain-Jones

## Abstract

Highly pathogenic avian influenza (HPAI) H5N1 clade 2.3.4.4b has expanded rapidly across the Southern Ocean since 2023, causing extensive mortality in sub-Antarctic wildlife. Yet its penetration into Antarctica and impacts on ice-dependent species remain poorly resolved primarily due to surveillance constraints. We report the first confirmed H5N1 infection in an Antarctic ice-dependent seal (crabeater seal; *Lobodon carcinophaga*) and document mortality of crabeater seals across the northern Weddell Sea during November–December 2024. Combining genomic, serological and observational data across nine species, we detected H5N1 RNA in a crabeater seal and a kelp gull (*Larus dominicanus*), and recovered complete HA, NA and M2 gene sequences from both. Phylogenetic analyses allowed us to identify at least two independent introductions of HPAI H5N1 clade 2.3.4.4b into the northern Antarctic Peninsula region. Serology provided strong evidence of prior exposure in scavenging birds, but no detectable H5 immunity in penguins or pinnipeds. Together, the results demonstrate ongoing novel viral incursions into Antarctica, likely facilitated by at-sea processes e.g. animal interactions on ice floes, that remain invisible to land-based surveillance. These findings highlight the vulnerability of ice-dependent pinnipeds to HPAI H5N1 clade 2.3.4.4b and the urgent need for expanded integrated Antarctic monitoring frameworks that pair serology, opportunistic carcass sampling and genomic sequencing.

## 1. Introduction

Since 2021, highly pathogenic avian influenza (HPAI) A/H5N1 clade 2.3.4.4b has expanded into a multi-continental panzootic affecting wildlife and domestic animals across the Americas, Europe, Africa, Asia and more recently the sub-Antarctic, with widespread mortality in birds and marine mammals (Krammer et al., 2025; Leguia et al., 2023). The first confirmed HPAI detections in the sub-Antarctic were brown skuas (*Stercorarius antarcticus*) at Bird Island, South Georgia, on 23 October 2023 (Bennison et al., 2024). In the Falkland Islands/Islas Malvinas, government testing confirmed infection in several seabird species from late October 2023 onward, with the number of cases suggesting local transmission (SCAR, 2024). Concurrent field observations and diagnostics documented unusual mortality in seals and several seabirds at South Georgia (e.g., skuas, gulls, penguins, shags) during this period (Banyard et al., 2024; Bennison et al., 2024).

By early 2024, HPAI had been confirmed south of 60°S on the Antarctic Peninsula. An intensive surveillance effort detected H5N1 clade 2.3.4.4b in dead skuas at James Ross Island (63.80°S, 57.81°W), constituting the first verified HPAI-associated mortality event on mainland Antarctica (Bennett-Laso et al., 2024). Subsequent genomic sequencing of brown skua viruses from the same area linked the Antarctic viruses phylogenetically to South American lineages, indicating at least one introduction from South America into Antarctica (Neira et al., 2025). Other contemporaneous reports included detections in kelp gulls (*Larus dominicanus*) from the South Shetland Islands (Ogrzewalska et al., 2024). The prior austral summer (2022/23) showed no evidence of HPAI in the northern Antarctic Peninsula and South Shetlands despite targeted surveillance (Muñoz et al., 2024). Collectively, these observations support a picture of delayed continental entry followed by rapid, localized spread once the virus breached the Peninsula (Lisovski et al., 2024).

Potential pathways for virus movement into, within, and out of Antarctica are consistent with known connectivity between South America and the Antarctic Peninsula via migratory seabirds, as previously demonstrated for low-pathogenicity avian influenza (LPAI) viruses and host movements (de Seixas et al., 2022; Hurt et al., 2016; Kopp et al., 2011). Ten months after the Antarctic mainland detections, HPAI outbreaks were confirmed on the French sub-Antarctic archipelagos of Crozet and Kerguelen, with mass southern elephant seal (*Mirounga leonina*) pup mortality documented (Clessin et al., 2025). Phylogeographic analyses indicated dispersal across the sub-Antarctic seeded from South Georgia (≈5,800 km to Crozet; ≈6,600 km to Kerguelen) rather than a single, purely local chain of transmission (Clessin et al., 2025). The elephant seal sequences from Crozet and Kerguelen are not monophyletic with the South Georgia sequence, therefore a single outward transmission wave from South Georgia to the French islands is unlikely. Instead, the topology indicates multiple novel introductions or the presence of unsampled ancestral lineages (Clessin et al., 2025).

H5N1 clade 2.3.4.4b has produced pinniped mortalities across multiple ocean basins, from mass die-offs of South American sea lions (*Otaria flavescens*) in Peru (Leguia et al., 2023) and elsewhere along the Pacific and Atlantic coasts (Plaza et al., 2024), to mortality in northern fur seals (*Callorhinus ursinus*) and Steller sea lions (*Eumetopias jubatus*) in the North Pacific (Sobolev et al., 2024), and European grey seals (*Halichoerus grypus*) in the North Atlantic (Mirolo et al., 2023), underscoring an unusually high susceptibility to the virus in marine mammals, to a nominally avian virus. Within the Antarctic region, H5N1 was confirmed in southern elephant seals and Antarctic fur seals (*Arctocephalus gazella*) on South Georgia during late 2023, with large-scale mortality reported at multiple sites (Bamford et al., 2025; Banyard et al., 2024). Because pinnipeds are long-lived, slow-reproducing apex consumers, sudden, large losses could propagate ecosystem effects and depress population trajectories for decades (Heithaus et al., 2008). The recovery of southern elephant seal populations is estimated to take until the end of the century (Campagna et al., 2025; Campagna et al., 2024), and continued losses may warrant reclassification of IUCN status. The transmission pathways into Antarctic ice-associated phocids remain unresolved: while predation or scavenging on infected birds may be implicated in other seal systems, obligate krill-specialists such as crabeater seals (*Lobodon carcinophaga*) rarely consume birds (Hückstädt et al., 2012), highlighting a key unknown in the potential spread of H5N1 in the Antarctic ecosystem.

Despite the scale of the panzootic, wildlife-focused surveillance in Antarctica remains poor due to logistical constraints: access is seasonal and weather-limited; diagnostic capacity is often off-continent and delayed; and carcass detection is imperfect in vast, remote landscapes (Wille et al., 2025), with the sea ice zone, that seasonally expands and contracts rapidly, representing a particular challenge. These constraints have motivated calls for innovative, opportunistic sampling networks integrating researchers, national programs, and the tourism sector (Wille et al., 2025) to improve sample collection across broader spatio-temporal scales.

Here, we leverage a multi-modal field program across the northern Antarctic Peninsula during November-December 2024. We (i) report the first viral genome from an Antarctic ice seal (a crabeater seal); (ii) document mortality in Antarctic ice seals; (iii) provide phylogenetic evidence for multiple incursions of H5N1 clade 2.3.4.4b into Antarctica; and (iv) demonstrate serological signatures of exposure and recovery in avian scavengers at mixed-species colonies. Together, these findings refine the pathways of HPAI establishment in the Antarctic region.

## 2. Methods

### 2.1 Study system

Fieldwork was carried out in December 2024 at three mixed-species breeding sites in the northern Antarctic Peninsula region (Figure 1). Two colonies—Brown Bluff (−63.533°, −56.917°) on the Tabarin Peninsula and the western shore of Joinville Island (−63.250°, −55.750°)—lie on the north-western margin of the Weddell Sea, whereas Penguin Island (−62.100°, −57.917°) is part of the South Shetland Islands archipelago in the Bransfield Strait. All sites support mixed assemblages of seabirds, including chinstrap, Adélie and gentoo penguins (*Pygoscelis antarcticus*, *P. adelie* and *P. papua*), scavenging birds (kelp gulls, brown skuas, snowy sheathbills *Chionis albus*) and hauled-out southern elephant seals and Antarctic fur seals on the rocky beaches, with hauled-out Weddell seals (*Leptonychotes weddellii*) and crabeater seals present on sea ice floes proximate to the landing sites. All colony landing sites appeared healthy at the time of landing, with no signs of unusual mortality. Across the three locations we obtained samples from 73 live individuals (60 birds, 13 pinnipeds) and 13 fresh carcasses spanning nine host species (Table 1).

**Figure 1.**
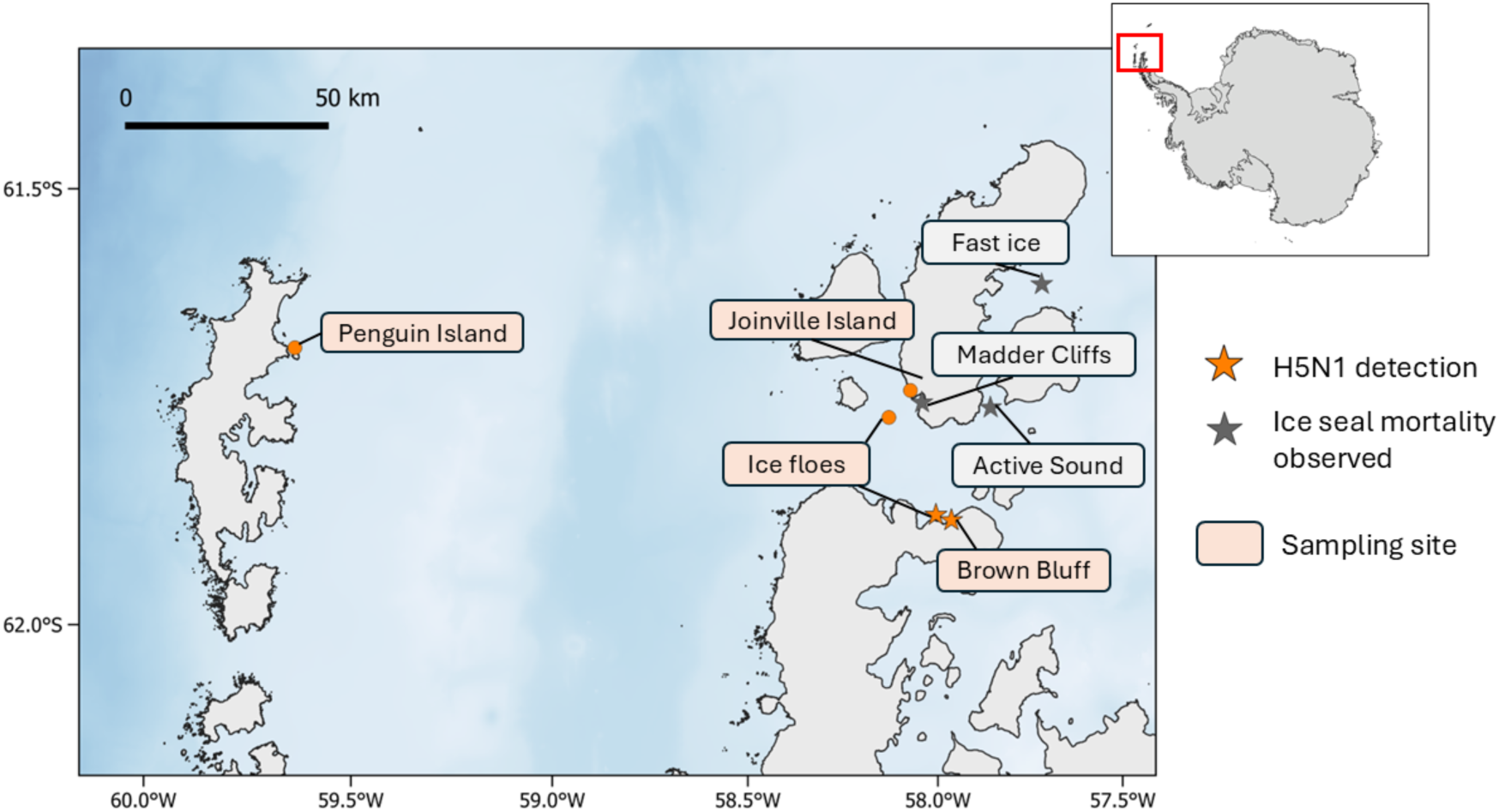
Sampling locations, confirmed H5N1 detections, and observed ice-seal mortality in the northern Antarctic Peninsula during November–December 2024. Map showing all sites surveyed during the expedition (orange circles), including Penguin Island, Joinville Island, Madder Cliffs, Active Sound and Brown Bluff, plus offshore sea ice floes. The location of confirmed H5N1 detections in a crabeater seal and a kelp gull are indicated with orange stars, and areas where ice-seal mortality was observed are marked with grey stars. The inset map shows the position of the study region relative to the Antarctic continent.

**Table 1.** Summary of avian influenza testing results from live seabird and pinniped species from northern Antarctic Peninsula.

| Location | Species | Sample size | H5 HPAIV +ve | AIV seropositive | H5 seropositive |
| --- | --- | --- | --- | --- | --- |
| Brown Bluff | Crabeater seal<br>(*Lobodon carcinophagus*) | 1 | 0 | 0 | 0 |
|  | Kelp gull<br>(*Larus dominicanus*) | 3 | 0 | 3 | 3 |
|  | Snowy sheathbill<br>(*Chionis albus*) | 5 | 0 | 5 | 3 |
|  | Brown skua<br>(*Stercorarius antarcticus*) | 1 | 0 | 1 | 1 |
|  | Adélie penguin<br>(*Pygoscelis adeliae*) | 15 | 0 | 2 | 0 |
|  | Gentoo penguin<br>(*Pygoscelis papua*) | 15 | 0 | 1 | 0 |
| Joinville Island | Weddell seal<br>(*Leptonychotes weddellii*) | 1 | 0 | 0 | 0 |
|  | Brown skua<br>(*Stercorarius antarcticus*) | 1 | 0 | 1 | 1 |
|  | Adélie penguin<br>(*Pygoscelis adeliae*) | 6 | 0 | 0 | 0 |
| Penguin Island | Southern elephant seal<br>(*Mirounga leonina*) | 8 | 0 | 0 | 0 |
|  | Brown skua<br>(*Stercorarius antarcticus*) | 3 | 0 | 3 | 3 |
|  | Chinstrap penguin<br>(*Pygoscelis antarcticus*) | 1 | 0 | 1 | 0 |

### 2.2 Ice seal mortality observations

Opportunistic observations of ice-seal carcasses were recorded onboard tourist vessels (*NG Endurance* and *NG Resolution*) in the north-western Weddell Sea and Antarctic Sound during November–December 2024. Observations were made opportunistically from the bridge and during sea ice landings, with carcasses identified and photographed on fast ice. Carcasses were tallied as minimum counts of clearly dead animals (“melting out” of the ice sensu visible submergence and fixed posture); condition (e.g. evidence of scavenging) were noted when discernible. We summarize these observations by date and location and treat them as lower-bound indicators of mortality extent rather than systematic surveys.

### 2.3 Sample collection

Sample collection from live animals consisted of oropharyngeal and cloacal swabs (birds), or nasal and rectal swabs (seals) using sterile nylon-flocked applicators. Swabs were stored in 1 mL of DNA/RNA Shield (Zymo Research) at −20°C until analysis. Whole blood was drawn from the brachial or metatarsal vein using a 1-mL heparinized syringe (birds) or extradural vein using a 5-mL heparinized syringe (seals). Red blood cells and plasma were separated by centrifugation and stored at −20°C until analysis. Samples were collected from carcasses opportunistically when encountered. Brain, oropharyngeal and cloacal swabs were taken and stored in 1 mL of DNA/RNA Shield (Zymo Research) at −20°C.

All sampling procedures were approved by the University of Tasmania Animal Ethics Committee (Project 31132). Fieldwork was conducted under permits issued by (i) the Australian Government Department of Climate Change, Energy, the Environment and Water, pursuant to the Antarctic Treaty (Environment Protection) Act 1980 (Permit 24/5243), and (ii) the US National Science Foundation (Permit ACA 2025-021).

### 2.4 Immunological assays

Plasma samples were evaluated for influenza-specific antibodies with two competitive ELISA assays from Innovative Diagnostics (France). The multispecies ID Screen® Influenza A Antibody Competition kit (FLUACA) detects antibodies to the highly conserved nucleoprotein (NP), whereas the ID Screen® Influenza H5 Antibody Competition kit (FLUACH5-V3) targets the H5 haemagglutinin. All procedures followed the manufacturer’s protocols. Optical densities (OD) were converted to percentage competition *(% competition*) using % 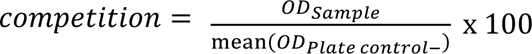. Because higher antibody concentrations yield lower OD readings, % competition declines as antibody levels rise. To present results intuitively, we report antibody binding as percent inhibition, calculated as 100 − % *competition*; negative inhibition values were set to zero, giving a 0 – 100 % scale (0 % = no detectable antibody, 100 % = strongest response). Samples were deemed seropositive when % inhibition met or exceeded the kit-specified cut-offs: ≥ 55 % for anti-AIV NP antibodies and ≥ 50 % for anti-H5 antibodies. Readings just below these thresholds (50 ≤. < 55 % for NP, 40 ≤. < 50 % for H5) are classified by the manufacturer as “doubtful.” To maintain conservative prevalence estimates, we treated these doubtful cases as negative. This affected only a small subset of samples (one for NP and one for H5); the vast majority fell well above or below the established cut-offs.

### 2.5 Virological Detection and Sequencing

Total RNA was extracted from swab samples using the Quick RNA Microprep kit (Zymo Research) following the manufacturer’s instructions. RNA was quantified using the Qubit^TM^ RNA HS Assay Kit (Thermo Fisher Scientific) on a Qubit^TM^ 4.0 Fluorometer (Thermo Fisher Scientific). Detection of the influenza A virus (IAV) followed a RT-qPCR assay protocol that targets a conserved region of the IAV matrix protein (MP) segment (Nagy et al., 2021). In brief, the assay was carried out using the QuantiNova Probe RT-PCR Kit (Qiagen), the primers SVIP-MP-F (5’-GGCCCCCTCAAAGCCGA-3’) and SVIP-MP-R (5’-CGTCTACGYTGCAGTCC-3’), and the TaqMan^TM^ MGB probe SVIP-MP_P2-MGB (5’FAM-TCACTKGGCACGGTGAGGGT-3’MGB-NFQ, Applied Biosystems^TM^). Thermocycling conditions were as per Nagy et al. (2021). Reactions were run on a Rotor-Gene Q (Qiagen) instrument using Rotor-Gene Q software version 2.3.5 (Qiagen). A Cq threshold of 36 was used to determine positive results.

RNA was reverse-transcribed and amplified with oligonucleotide primers designed to amplify all eight viral segments simultaneously (Hoffmann et al., 2001; Zhong et al., 2019), in a single step RT-PCR using the SuperScript III One-Step RT-PCR kit (Thermo Fisher Scientific). PCR products were purified using AMPure XP Beads (Beckman Coulter) at 1.8x. Library preparation was conducted using the Rapid Barcoding Kit 24 V14 (Oxford Nanopore Technologies) following the manufacturer’s protocol, and libraries were sequenced on a PromethION 2 Solo unit using a PromethION flow cell (R10.4.1 chemistry, Oxford Nanopore Technologies).

To fill in sequencing gaps, shotgun RNA sequencing libraries were prepared by 3’ polyadenylation of the extracted RNA using the E. coli Poly(A) Polymerase kit (New England Biolabs), followed by library preparation with the cDNA-PCR Sequencing V14 – Barcoding kit (Oxford Nanopore Technologies) following the manufacturer’s protocol. Sequencing was performed on a MinION flow cell (R10.4.1 chemistry, Oxford Nanopore Technologies) with a MinION device.

### 2.6 Bioinformatics and phylogenetic reconstruction

Basecalling was performed after the sequencing run on the MinKNOW software (Oxford Nanopore Technologies), using the super high accuracy setting. Trimmed super high accuracy reads were mapped to the H5N1 segments using Minimap2 (Li, 2018) with default ONT parameters. To address low coverage in some AIV segments, reads from both PCR-amplified and shotgun libraries were pooled, yielding a minimum of 10× coverage (often exceeding ∼10,000×). Consensus HA sequences were then aligned in Geneious Prime version 2025.0.2 (Kearse et al., 2012) using the Clustal Omega algorithm (Sievers & Higgins, 2013) against 331 background HA sequences (all wildlife 2.3.4.4b, avian excluding poultry and mammals, from South America and South Africa) downloaded from GISAID on 7 June 2025. The same pipeline was applied to the M2 and NA regions to confirm introduction patterns.

For each gene, maximum-likelihood phylogenies were inferred in IQ-TREE 2 (Minh et al., 2020) with 1,000 ultrafast bootstrap replicates (Hoang et al., 2018) and substitution models selected via ModelFinder (Kalyaanamoorthy et al., 2017). Final consensus trees were visualized in R using ggtree (Yu et al., 2017).

## 3. Results

### 3.1 Ice seal mortality observations

Opportunistic ship-based observations documented unusual crabeater seal mortality on fast ice around the northern Weddell Sea in late spring 2024 (Figure 2). On 15 November 2024 in Active Sound (−63.440°, −56.339°), 37 crabeater seals were recorded dead within ∼0.5 nautical miles of the ice edge; carcasses appeared to have been dead for some time and were clustered along tidal cracks, “melting out” of the ice. On 28 November 2024, one additional carcass was observed near Seymour Island. On 6 December 2024 near Madder Cliffs (−63.3069°, −56.5064°), 4 dead adult crabeater seals were observed ∼1.5 nautical miles from the ice edge with initial scavenging evident. Finally, on 25 December 2024 observers reported hundreds of dead seals on fast ice in Antarctic Sound (−63.3812°, −55.6897°). Taken together, these minimum counts indicate that the crabeater seal carcass we sampled formed part of a broader mortality event on fast ice preceding seasonal break-out.

**Figure 2.**
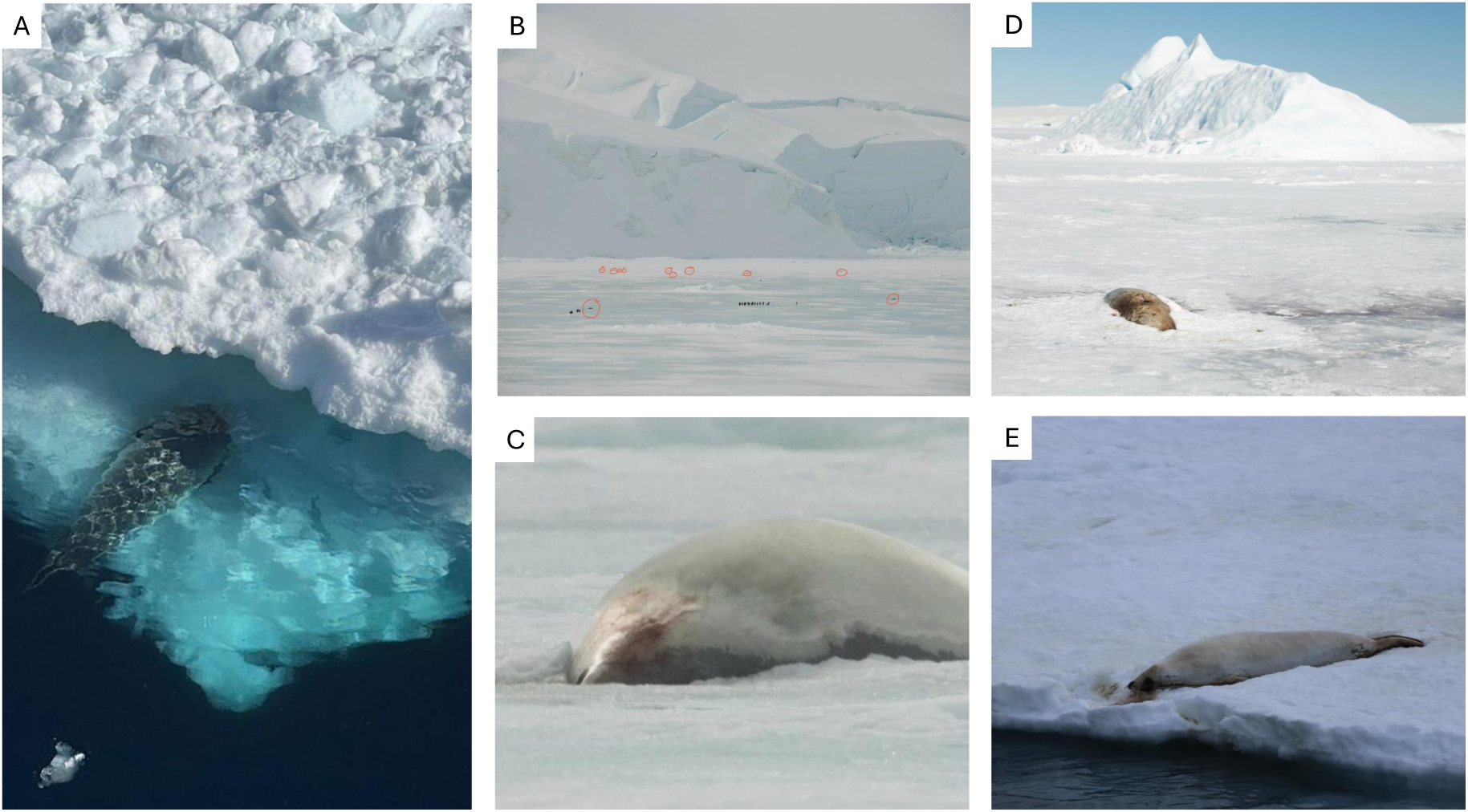
Opportunistic observations of crabeater seal mortality on fast ice, Weddell Sea (Nov–Dec 2024). (A) 28^th^ November, near Seymour Island, one seal; (B-C) 15^th^ November, Active Sound, 37 crabeater seals; (D) 6^th^ December, Madder Cliffs, four crabeater seals; (E) 25^th^ December, “hundreds of dead seals on fast ice”. Image A) ©Jonathan Reid, all other images ©Jessica Farrer.

### 3.2 Past exposure to H5Nx of Antarctic wildlife

Both AIV and H5 seroprevalences differed markedly among foraging guilds. Scavenging species exhibited clear evidence of prior exposure to H5Nx. All tested kelp gulls, brown skuas, and snowy sheathbills had AIV antibodies, and all kelp gulls and brown skuas, and 60% of sheathbills, were positive for H5-specific antibodies (Tables 1–2). H5-seropositive birds were found at all sampled sites. These results indicate that scavenging birds at mixed-species colonies had encountered H5Nx and mounted an immune response, consistent with their frequent consumption of marine mammal and seabird carrion. In contrast, all sampled pinnipeds were seronegative for both AIV and H5-specific antibodies (Table 2). The absence of detectable immunity in elephant seals is notable given the extensive seal mortality reported at sub-Antarctic sites, and likely reflects the demographic composition of the Antarctic cohort: the individuals sampled in this study were young, pre-breeding animals that haul out in the Weddell Sea rather than at the high density sub-Antarctic breeding colonies where exposure risk is higher. None of the 37 penguins tested positive for H5 antibodies (Tables 1–2). AIV antibodies were detected in a small number of individuals.

**Table 2.** Summary of H5 seroprevalence of seabird and pinniped species from northern Antarctic Peninsula.

| Species | Sample size | H5 Seroprevalence (%) |
| --- | --- | --- |
| Crabeater seal<br>(*Lobodon carcinophagus*) | 1 | 0.00 |
| Weddell seal<br>(*Leptonychotes weddellii*) | 1 | 0.00 |
| Southern elephant seal<br>(*Mirounga leonina*) | 8 | 0.00 |
| Kelp gull<br>(*Larus dominicanus*) | 1 | 100 |
| Snowy sheathbill<br>(*Chionis albus*) | 5 | 60.0 |
| Brown skua<br>(*Stercorarius antarcticus*) | 5 | 100 |
| Gentoo penguin<br>(*Pygoscelis papua*) | 15 | 0.00 |
| Adélie penguin<br>(*Pygoscelis adeliae*) | 21 | 0.00 |
| Chinstrap penguin<br>(*Pygoscelis antarcticus*) | 1 | 0.00 |

**Table 3.**
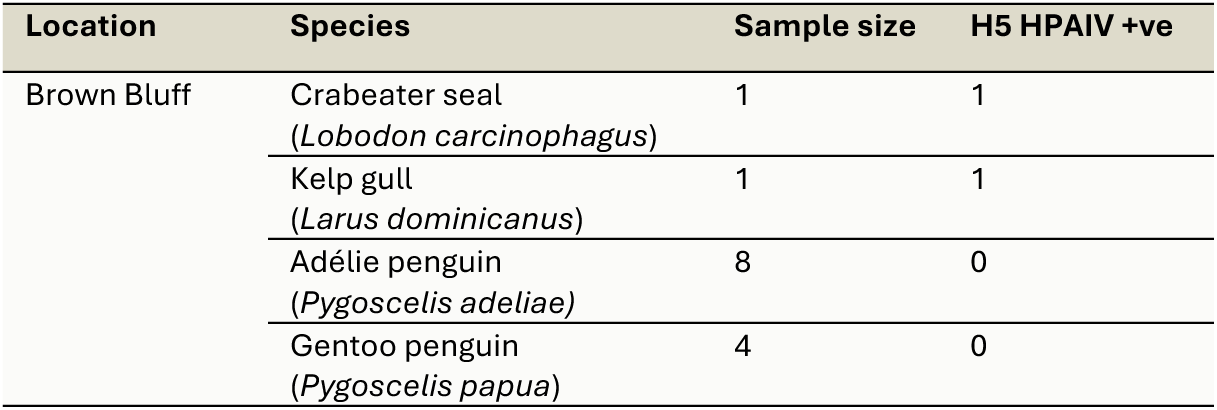
Summary of avian influenza testing results from carcasses of seabird and pinniped species from northern Antarctic Peninsula.

### 3.3 H5N1 clade 2.3.4.4b in Antarctic wildlife

H5N1 clade 2.3.4.4b was detected in two carcasses: a crabeater seal and a kelp gull. All other individuals tested negative by RT-qPCR. Complete HA, NA and M2 gene segments were obtained from both the crabeater seal and kelp gull carcasses (Genbank Accessions PX632568-PX632573). Maximum-likelihood phylogenetic analysis placed both sequences within clade 2.3.4.4b (Figure. 3), clustering among wildlife-derived South American lineages. The two Antarctic sequences did not form a monophyletic group; instead, each fell into a distinct sub-lineage. These phylogenetic positions indicate at least two independent introductions of H5N1 into the northern Antarctic Peninsula region. The crabeater seal sequence was most closely related to lineages circulating in South American pinnipeds and seabirds, whereas the kelp gull sequence fell within a separate cluster associated with Patagonia and the Falkland Islands.

**Figure 3.**
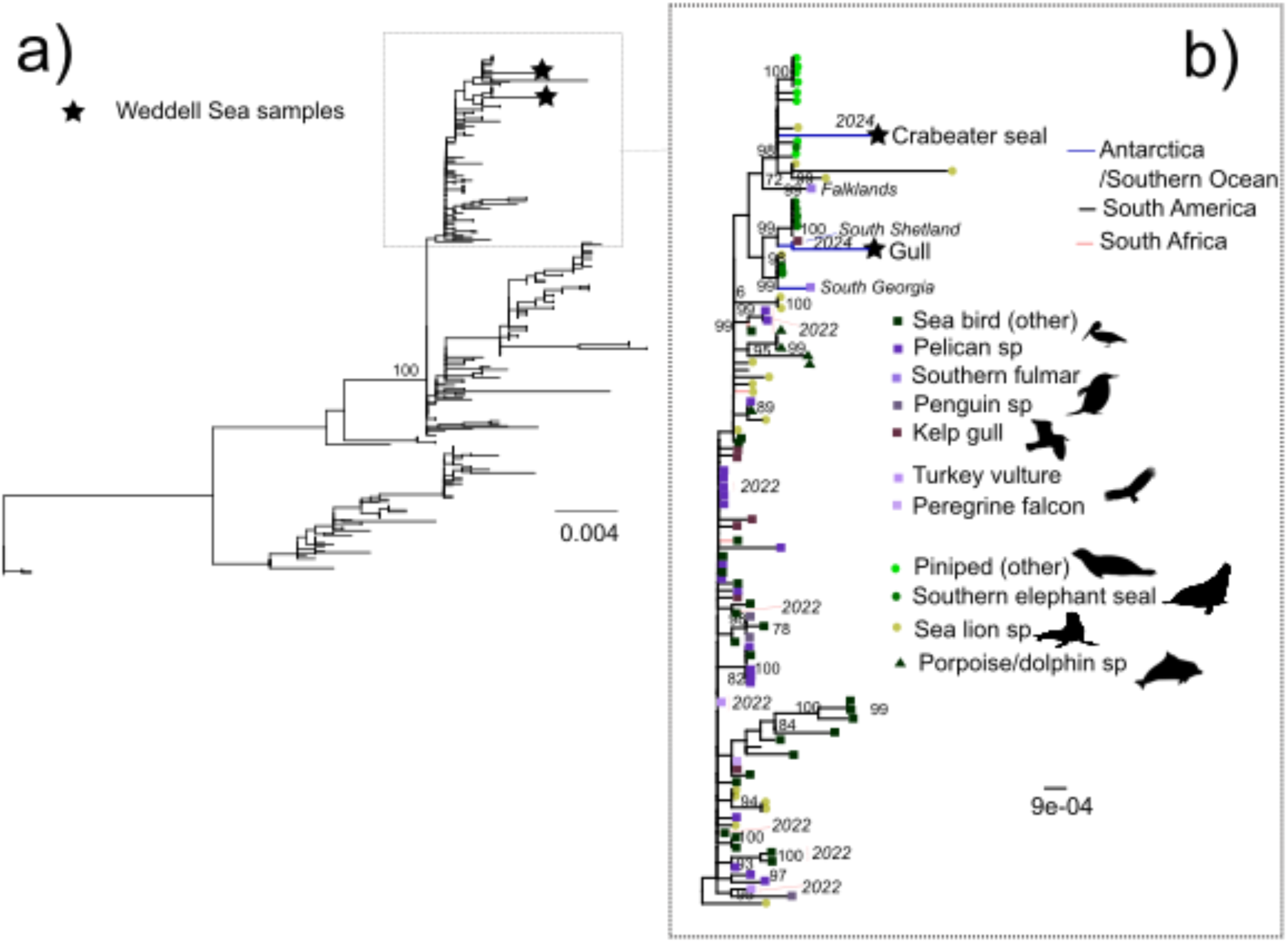
Maximum-likelihood phylogeny for the HA gene depicting the placement of our Wedell Sea crabeater seal and gull HPAI 2.3.4.4b sequences within multi-species circulation across South America, South Africa and the sub-Antarctic. Tip symbols denote host origin—squares for avian, circles for pinniped and triangles for other marine mammals—and are colour-coded accordingly. Node labels indicate ultrafast bootstrap support values. Panel (A) shows the full tree containing 333 sequences; inset (B) highlights the major clade encompassing our study samples.

## 4. Discussion

Our findings confirm, for the first time, infection of ice seals with HPAI H5N1 and reveal the complex, multi-host epidemiological landscape of this virus in the Southern Ocean. By integrating genomic data, serology and ecological context, we show that the virus entered the northern Weddell Sea via at least two independent introduction events, consistent with patterns of sub-Antarctic dispersal documented across the wider Southern Ocean during 2023–2025 (Clessin et al., 2025; Kuiken et al., 2025; Leguia et al., 2023; Lisovski et al., 2024; Neira et al., 2025). In addition, detection of H5-seropositive individuals are all sampled sites suggest a wide circulation of the virus and/or potential carriers across the region. Although the sub-Antarctic islands have experienced large-scale mortality, particularly of pinnipeds (Bamford et al., 2025; Clessin et al., 2025), extensive population-level die-offs have not yet been observed in Antarctica, despite evidence of viral incursion into multiple species and sites. This raises an important question: does this apparent disparity reflect genuine differences in animal behaviour and landscape structure that shape disease outcomes in Antarctic wildlife compared to sub-Antarctic wildlife, or have large-scale mortality events simply gone undetected across the vast and challenging (difficult to observe) sea-ice environment?

### 4.1 Introductions and connectivity with the broader Southern Ocean

Our phylogenetic analyses support a scenario in which the northern Weddell Sea region was seeded from South America via multiple, distinct introductions, mirroring the dominant eastward trans-basin movement of H5N1 across the sub-Antarctic (Clessin et al., 2025; Lisovski et al., 2024). These regional patterns reflect the repeated dispersal of the virus from South America into South Georgia, the Falkland Islands and, subsequently, south into the Antarctic Peninsula, as well as eastward over > 6,000 km to the southern Indian Ocean islands of Crozet and Kerguelen – movements facilitated by the long-distance foraging and migratory behaviour of albatrosses, giant petrels and skuas (Aguado et al., 2024; Clessin et al., 2025; León et al., 2025; Lisovski et al., 2024). The strong eastward transmission bias observed throughout the Southern Ocean aligns with prevailing wind systems and the circumpolar distributions of wide-ranging seabirds, which regularly traverse ocean basins (Hindell et al., 2020; Lisovski et al., 2024).

The repeated movement of H5N1 across vast expanses of open ocean demonstrates that long-distance dispersal is not a one-off anomaly but an inherent feature of the current panzootic. These events may be facilitated through at-sea processes that remain invisible to land-based monitoring, including the scavenging of floating pinniped and seabird carcasses by highly mobile scavengers such as skuas and giant petrels (Lisovski et al., 2024). Offshore transmission pathways also offer a plausible mechanism for viral introduction into crabeater seals, which occupy extensive pack-ice habitats and lack long-range migratory connectivity with sub-Antarctic breeding sites (Hückstädt et al., 2020). In crabeater seals, exposure may occur through marine interfaces, for example, via indirect contact with infected seabirds on shared sea ice floes i.e. infection via environmental exposure, rather than via direct transmission from sub-Antarctic pinniped species. This highlights the importance of considering offshore epidemiological processes when interpreting patterns of Antarctic infection.

### 4.2 Ice seal infection detection challenges

The unique ecology of Antarctic ice-dependent pinnipeds makes mortality exceptionally difficult to detect. Unlike Antarctic fur seals and southern elephant seals, which haul out on accessible beaches where carcasses accumulate and can be readily counted (Bamford et al., 2025; Clessin et al., 2025), crabeater, Weddell, leopard, and Ross seals live primarily on sea ice. The sea ice zone is a dynamic, fragmented habitat where carcasses are rapidly obscured by snow, redistributed by wind, or lost as ice breaks out and melts seasonally. Unlike beaches or sub-Antarctic haul-outs, the pack-ice environment spans millions of square kilometres and is almost entirely inaccessible to observers. As such, detection probability for dead ice seals is low and large-scale mortality events could easily go undetected. In this context, the few carcasses documented here should be interpreted as the visible fraction of a potentially much larger event. In the absence of additional H5N1 sequences from ice seals, it is unknown whether this wider mortality represents a monophyletic clade, and therefore a single rare event of spillover to crabeater seals, or if there have been multiple transmissions to ice seals from other taxa. There is therefore an urgent need for additional live and carcass sample of ice seals, plus ongoing monitoring of ice seal population sizes, for example via high-resolution satellite imagery (LaRue et al., 2022), to quantify potential impacts on ice-dependent pinnipeds.

### 4.3 Species-specific responses and the ecology of exposure

The northern Weddell Sea community shows a clear, species-specific pattern of exposure. Notably, all southern elephant seals we sampled tested negative by both PCR and serology, despite the widespread H5N1-associated mortality documented for the species across the sub-Antarctic (Bamford et al., 2025; Clessin et al., 2025). Most elephant seals present on the north-eastern Antarctic Peninsula beaches, during the survey period, are juveniles. Their more dispersed haul-out behaviour and reduced connectivity to dense breeding colonies could limit opportunities for transmission, in contrast to the tightly aggregated pupping and moulting colonies that experienced mass mortality in the sub-Antarctic.

Scavenging birds showed clear evidence of exposure. We detected one active infection in a kelp gull and identified multiple seropositive individuals across kelp gulls, skuas and snowy sheathbills. Their foraging ecology includes regularly feeding on marine mammal and bird carcasses, providing a direct interface with infectious material. The presence of anti-H5 antibodies in several individuals indicates that, despite early records of massive mortality events affecting notably skuas (Bennett-Laso et al., 2024), these species are surviving infection and may now possess at least partial immunity, with overall high seroprevalence among scavenging taxa. Although our limited sampling does not allow us to assess whether these taxa support sustained intraspecific transmission, the serological patterns suggest that scavengers may be repeatedly exposed and capable of mounting effective immune responses, reinforcing their utility as ecological sentinels for early detection of viral incursions (Lisovski et al., 2024).

By contrast, penguins showed no evidence of H5Nx infection, with all individuals testing negative by PCR and serology. Given the presence of the virus in other taxa at the same colony sites, the absence of infection in penguins suggests either that they have lower exposure risk than neighbouring skuas, sheathbills and kelp gulls, or that infection is highly lethal in penguin species, with exposed individuals dying rapidly and failing to develop detectable antibody responses. Their pelagic foraging ecology and lack of scavenging behaviour likely reduce opportunities for carcass-mediated transmission, which appears to be a major pathway in Antarctic and sub-Antarctic systems. Overall, these patterns indicate that exposure risk is tightly linked to foraging ecology.

Overall, the Weddell Sea assemblage shows a pattern of multiple introductions and limited transmission, with infection confirmed in a crabeater seal and a kelp gull, and additional exposure detected in several scavenging species. Despite this, we found no evidence of amplification into colony-level outbreaks, and key taxa such as penguins, southern elephant seals, and Weddell seals remained uninfected in our dataset. This contrasts with the severe, rapid-onset mortality events documented at sub-Antarctic breeding sites (Bamford et al., 2025; Clessin et al., 2025). Together, these findings underscore that in Antarctica, H5N1 dynamics are shaped by the ecological context: diffuse, ice-associated haul-outs, low-density interactions, and foraging ecology-linked transmission. The species driving the spread of H5N1 in Antarctic remain unclear and further sampling of both live animals and carcasses is needed to understand transmission dynamics in the region.

### 4.4 Implications for future outbreaks and endemisation potential

A central question is whether H5N1 could become seasonally maintained in the Weddell Sea. Evidence from sub-Antarctic islands shows the virus can overwinter in wildlife communities (Lisovski et al., 2024), raising concerns about endemic dynamics. Several features of the Weddell system may limit persistence: predator and scavenger assemblages are less diverse, colony structure is more fragmented, and winter access to pinniped hosts is minimal. However, repeated long-range transmission events are a key feature of the ongoing panzootic, and future outbreaks may differ substantially from the low-intensity spillovers observed here.

While our genomic data are insufficient to confirm mammal-adapted transmission in Antarctica, PB2 mammalian adaptations detected elsewhere (Kuiken et al., 2025; Neira et al., 2025) underscore the potential for onward adaptation. The observation of large numbers of clustered seal carcasses reported here suggests seal-to-seal transmission is likely occurring in Antarctic ice-dependent pinnipeds.

### 4.5 Conclusion and surveillance implications

Our findings show that HPAI H5N1 has now reached ice-dependent Antarctic pinnipeds and is circulating within a uniquely challenging Antarctic environment where transmission pathways, host behaviour and detection probability differ sharply from those in the sub-Antarctic. The Antarctic system appears to experience low-amplification introductions, driven largely by scavenging-mediated exposure and at-sea processes, rather than the explosive, colony-level outbreaks documented elsewhere. This underscores the need for surveillance approaches that move beyond carcass-based detection.

Across the Southern Ocean, three lessons are becoming increasingly clear. First, PCR-only surveillance substantially underestimates outbreak extent, particularly where transmission likely occurs offshore and carcasses are rapidly lost in sea-ice habitats. Second, serological sampling of apex predators and scavengers provides an early, spatially integrative indicator of viral circulation. Third, genomic sequencing, even from a handful of carcasses, can reveal hidden connectivity and introduction routes. Given the severe logistical constraints of Antarctic fieldwork, an integrated surveillance framework that combines serology, opportunistic carcass sampling and targeted genomic sequencing will be essential for detecting introductions early, understanding how the virus is moving through the system, and anticipating future outbreaks. As H5N1 continues to evolve and expand its ecological footprint, proactive, multi-layered monitoring is critical to understand the broader implications of this panzootic for Southern Ocean ecosystems.

## Acknowledgements

This research was supported by the National Geographic and Rolex Perpetual Planet Ocean Expeditions and conducted in collaboration with the Schmidt Ocean Institute aboard the *R/V Falkor (too)* (Cruise ID FKt251214). We acknowledge funding from the Geoffrey Evans Trust, Kenneth C. Griffin, and Griffin Catalyst. The authors would like to thank Elizabeth Hogan, Sujatha Bagal, Nik Varley, Nicole Alexiev, Tyler Dinley, Maria Sanchez and Veit Huehnerbach, for facilitating sample collection and their ongoing support of this research.

## Author Contributions

**Jane Younger**: Conceptualization (lead); writing – original draft (lead); investigation (equal); formal analysis (equal); writing – review and editing (equal); funding acquisition (lead); project administration (lead). **Amandine Gamble**: Conceptualization (equal); investigation (equal); writing – review and editing (equal); funding acquisition (supporting); project administration (supporting). **Nicholas Fountain-Jones**: Formal analysis (lead); visualization (lead); writing – original draft (supporting); investigation (equal); writing – review and editing (equal). **Laura Patier**: writing – original draft (supporting); investigation (supporting); writing – review and editing (equal). **Talia Brav-CubiW**: writing – original draft (supporting); investigation (supporting); writing – review and editing (equal). **André van Tonder**: investigation (supporting); writing – review and editing (equal). **Augustin Clessin**: investigation (supporting); writing – review and editing (equal). **Jamie Coleman**: investigation (supporting); writing – review and editing (equal). **Jessica Farrer**: investigation (supporting); writing – review and editing (equal). **Clive McMahon**: Conceptualization (supporting); investigation (equal); writing – review and editing (equal); funding acquisition (supporting).

