## Supplemental Figures for "High pathogenicity avian influenza virus H5N1 clade 2.3.4.4b in Antarctica: Multiple Introductions and the First Confirmed Infection of Ice-Dependent Seals"

Supplementary Figures

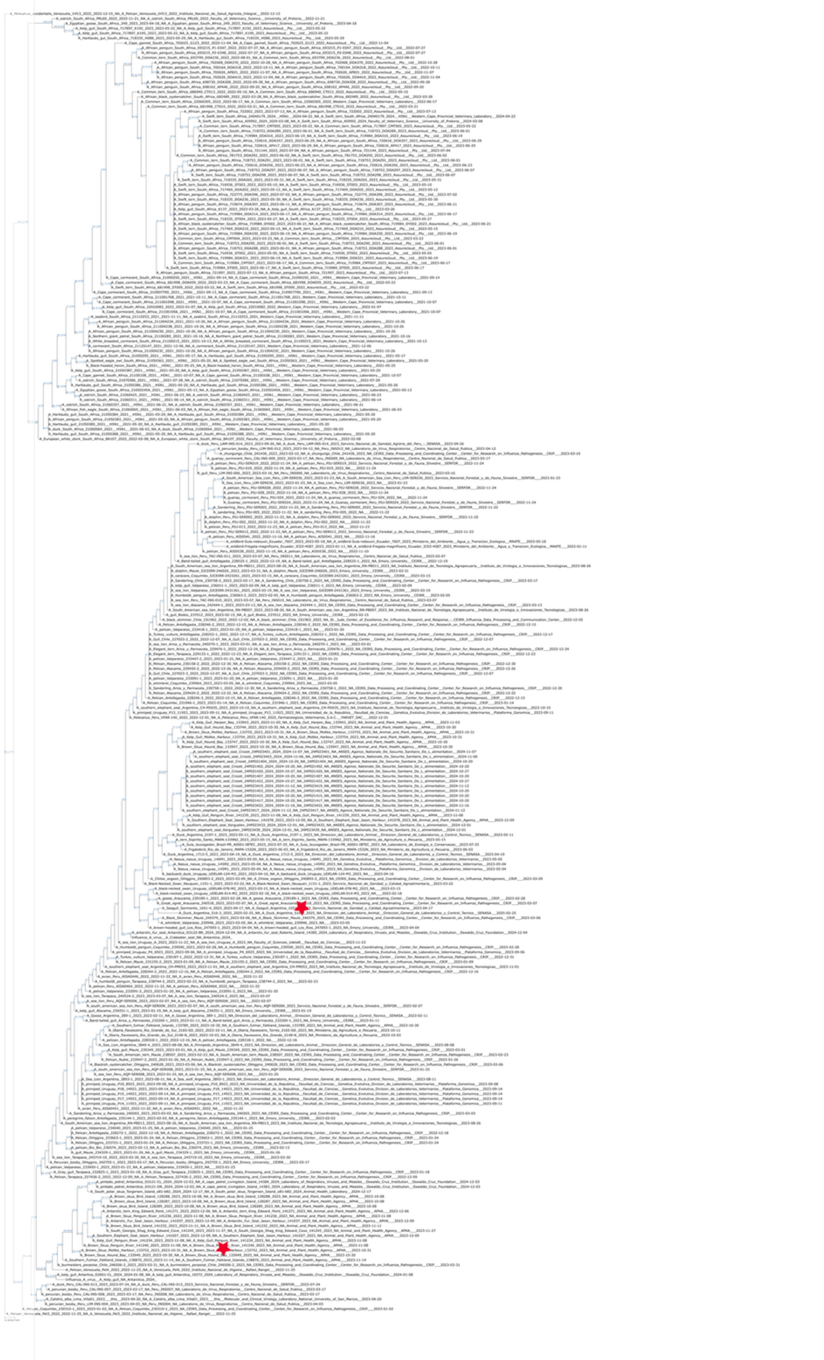

**Figure S1**. Maximum-likelihood phylogeny for the NA gene depicting the placement of our Wedell Sea crabeater seal and gull HPAI 2.3.4.4b sequences (indicated with red stars) within multi-species circulation across South America, South Africa and the sub-Antarctic. Node labels indicate ultrafast bootstrap support values.

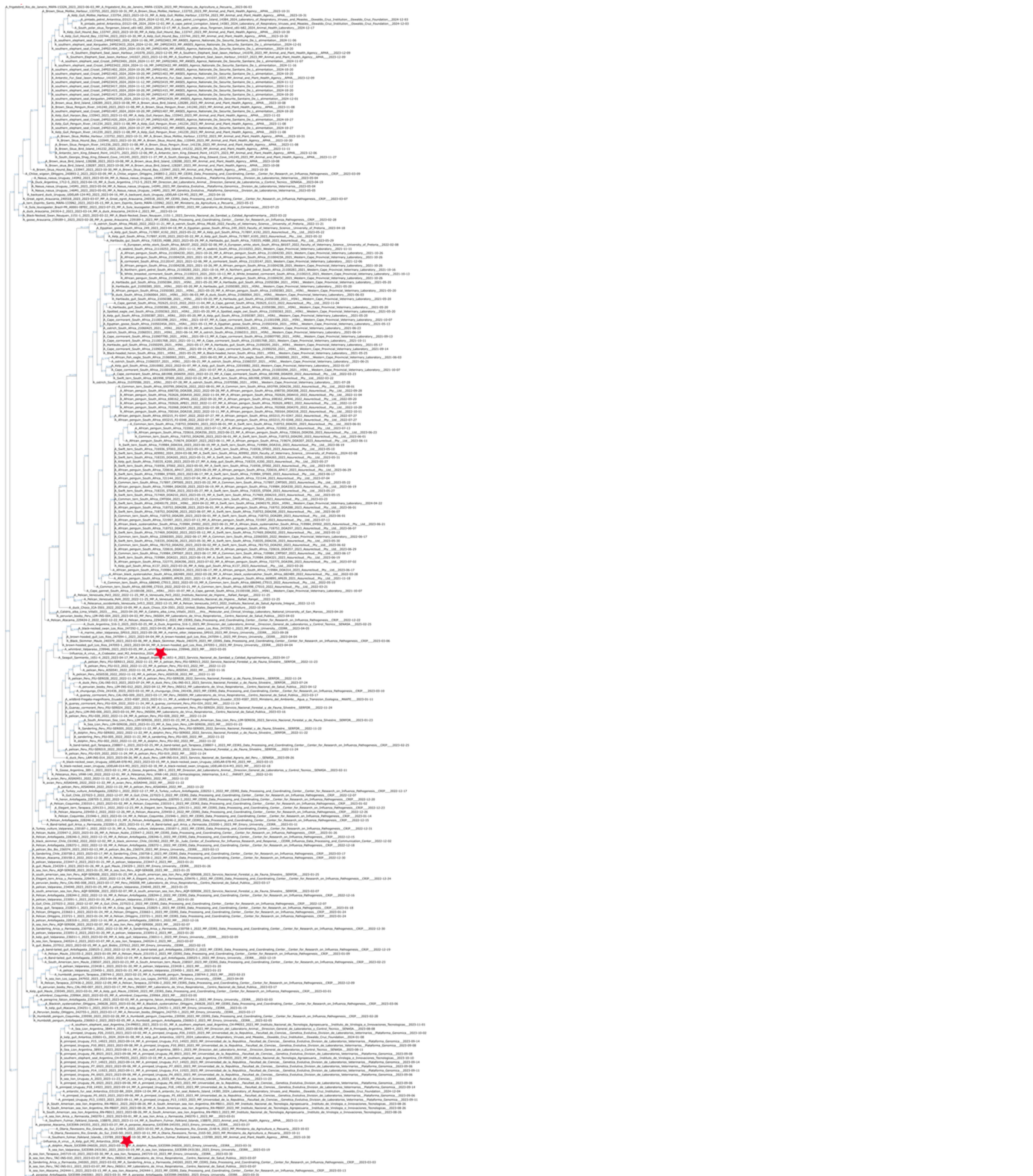

**Figure S2**. Maximum-likelihood phylogeny for the M2 gene depicting the placement of our Wedell Sea crabeater seal and gull HPAI 2.3.4.4b sequences (indicated with red stars) within multi-species circulation across South America, South Africa and the sub-Antarctic. Node labels indicate ultrafast bootstrap support values.
